# Drug delivery via tattooing: Effect of needle and fluid properties

**DOI:** 10.1101/2021.02.02.429454

**Authors:** Idera Lawal, Pankaj Rohilla, Jeremy Marston

**Author notes:** Corresponding author ✉.

## Abstract

Tattooing is a commonplace practice among the general populace in which ink is deposited within dermal tissue. Typically, an array of needles punctures the skin which facilitates the delivery of a fluid within the dermis. Although, a few studies in the past have investigated the potential of tattooing as an intradermal (ID) drug injection technique, an understanding of the fluid dynamics involved in the delivery of fluid into skin is still lacking. Herein, we sought to provide insight into the process via an in vitro study. We utilize a five needle flat array (5F) with a tattoo machine to inject fluids into gelatin gels. High-speed imaging was used to visualize the injection process and estimate the amount of fluid delive red after each injection upto the 50^th^ injection. We investigate the role of reciprocating frequency (*f*) of the needle array and the physical properties of the fluids on the volume (V_o_) and the percentage delivery (*η*) after injection. In addition, we illustrate the physical mechanism of fluid infusion during tattooing, which has not been reported. An understanding of the injection process via tattooing can be useful in the development of ID tattoo injectors as drug delivery devices.

## INTRODUCTION

The tattooing technique reportedly dates back to the Neolithic period from ancient artifacts discovered by archaeologists [1], but only to the fourth millennium BC based on tattoos found on mummified human skin [2]. The artifacts and skin samples suggest that the tattoos were either intended for cosmetic aesthetic or for the concealing of body defects such as baldness, scars and the appearance of a after mastectomy [3].

To date, tattooing is widely practiced and the number of people with at least one tattoo has seen major increase worldwide [2]. An estimated 1 in 3 Americans have at least one tattoo with other countries like Italy or Sweden having even higher statistics [4]. Several studies have shown, that skin type and individual innate immune system make up are key factors in whether the tattoo results in allergic reactions and pain, these factors also determine how much of the ink is injected (20% – 50% effective volume) [5]. Therefore, from population magnitude alone, the process poses motivation for scientific and technological inquiry into abating health risk and improving patient compliance.

The tattooing procedure involves the invasive repeated puncture by 200-350 micron sized needles, intradermally (0.5-2 mm depth). This effectively injures the topmost part of the dermis as the needles drive more ink in with each successive puncture. Needles used vary in both number and conformation, and the amount of ink delivered is dependent on ink/skin properties, the depth and density of punctures. Tattoo inks are colorants made of metallic salts which are suspended in a carrier solution along with binders, surfactants and other additives [5]. The specific composition of the inks is kept secret by manufacturers, however because each formulation is tailored to optimum fluid dynamic properties for delivery, rheological behavior suffices for characterization [4]. The number of needles range from the single needle head to some 23 needles.

After the procedure, the body immune system kicks into overdrive (from the thousands of micro injuries that just occurred) and responds by; inflammation which is immediately triggered by tissue phagocytes at the scene of injury. These cells release histamines that cause a transient increase in vasodilation and vascular permeability accompanied with the release of cytokines that signal the next phase; influx of monocytes through the blood vessels (aided by vasodilation). Monocytes become macrophages when they reach the site of injury, after which they either consume the deposited ink and traverse to the lymph nodes or remain in the dermis (a reason for the seemingly permanent nature of tattoos) [2].

DNA vaccination and some novel therapeutic treatments also involve intradermal delivery. Over the past century, DNA vaccination has evolved from volume profligate intramuscular injections to antigen – sparing intradermal doses that have similar or improved immune responses [6]. Several studies have shown that this simple delivery route change enhanced immunogenicity and thus reduced cost [7, 8, 9, 10]. This Fractional dosing involved subjects receiving a single 20% shot of the full dose. However, challenges such as needle-stick injuries and cross contamination persist. Jet injectors [6]; micro-needle patches [11]; ballistic vaccination methods such as the gene gun [12] and epidermal powder immunization [13]; are examples of needle free alternatives that circumvent such difficulty. Moreover, the delivery of DNA vaccines can also be challenging. It constitutes deeply entangled, high molecular weight, highly viscous shear thinning suspensions which is sensitive to hydrodynamic shear [14]. For this reason, the needle free jet injector is preferred due to the high shearing that occurs *O*(~ 10^6^*_s_*^-1^) albeit concerns about high shear degradation have been reported [14].

The extent to which the immune system responds has driven several investigations into the efficacy of the tattooing process in intradermal (ID) drug delivery [5]. Some studies adapt the tattoo machine for even injection into a large area, effectively portioning medication into fractional doses [15]. To state a few, Shio et al. [16] utilize a 5 needle magnum head for the treatment of cutaneous leishmaniasis in their study. Each session consisted of a fractional dosage of 12 two-second applications twice everyday for five days, successfully administering approximately 2-5 microlitres of treatment to attenuate the effects of the disease. In a similar study, Quaak et al. [15] conduct in vitro and ex vivo tattooing of plasmid DNA solutions using a 9 needle flat head to probe the extent to which their Murine models had antigen expression and T-cell responses. Furthermore, Potthoff et al. [17] confirmed that in comparison to ID injections, tattoo immunization restrict antigen expression to the topmost layer of the dermis even though expression levels are similar. In this study, an 11 magnum head needle is used for 10 microliter fractional doses. Studies on the tattooing process are not restricted to vaccination and treatment of disease as probe into the immune response [18] and recovery time for tattoo inks in a bid to illuminate on the toxicology of tattoo inks and pigments. To do this, they utilize a 14 magnum needle head. Interestingly, the basis for which the number of needles used is not usually reported in these studies, and because majority of these researchers stem from toxicology, immunology and applied pharmacology backgrounds, it is reasonable to suggest that the choice is arbitrary.

As far as the physics of the process, very few literature exists that detail the intricacies of the process [19, 20] and more rigorous descriptions of the physics behind the tattooing procedure is lacking despite the wealth of literature on the immunological response of the process.

In this study, we use a transparent gelatin gel to understand the injection mechanism via tattooing process. We used DI water, 80% glycerol as Newtonian fluids and two commercial inks (black and red ink) exhibiting non-Newtonian behavior as fluids to be injected using a tattoo pen equipped with a five needle array in a flat conformation. We compared the volume delivered for different number of repeat injections and varying needle reciprocating frequency.

## MATERIALS AND METHODS

An atom pen rotary tattoo machine (*Dragonhawk Tattoo Supply, China*) was used. A 12-gauge 5F tattoo needle (*five needles arranged in a flat array*) with each needle having a total length of 10 mm (tapered length ~1 mm from the tip) and a diameter of 0.35 mm, was used with the tattoo machine. DI water and 80% glycerol (G,*Macron*) were used in addition to the black and red ink pigments (*Mom’s Millenium, USA*) as injectates. The reciprocating frequency of needles (46-72 Hz) was varied via voltage control in a range of 3-5 V. Figure 1(a) shows the schematics of the experimental setup used.

**Figure 1:**
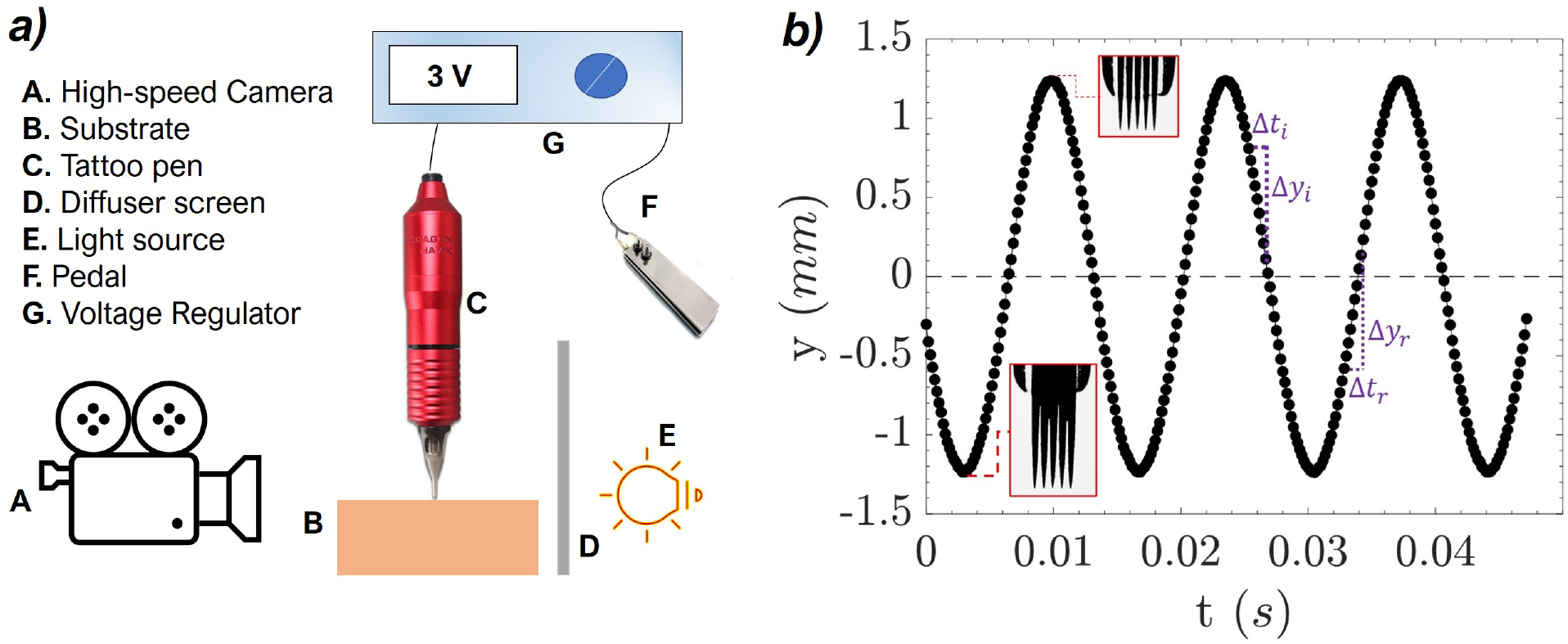
Experimental. **(a)** Schematics of experimental setup and (**b)** needle displacement frequency of a 5F tattoo needle for a voltage of 3 V.

Rheological measurements of ink pigments and 80% glycerol were performed on a DHR-2 rheometer (TA *Instruments, USA*) with a cone and plate geometry (25 mm diameter and 1.992° angle) and at a temperature of 22±1 °C. The flow behavior of ink pigments and 80%G was measured via flow ramp tests for applied shear rate ranging from 0.001-5000 s^-1^, whose results are presented in the Appendix.

A high-speed camera (*Phantom V711, Vision Research Ltd*) was used to capture the motion of the reciprocating needles and injections at a frame rate of 8000 fps with pixel size of 16μm/px (Spatial resolution: 400×800). A Nikon micro-nikkor 60mm lens was used with a 28mm extension tube. Figure 1(b) shows the reciprocating motion of the needle with a frequency of 46 Hz corresponding to a voltage of 3 V. Insertion velocity (*υ*_i_ = Δ*y_i_*/Δ*t_i_*) and retraction velocity (*υ*_r_ = Δ*y_r_*/Δ*t_r_*) were estimated by tracking the needle tip using image processing via a custom Matlab Script. It is noteworthy that these velocities do not deviate much even inside the substrate.

An in vitro study was conducted to understand the mechanism of drug delivery via tattooing. Gelatin powder (*from bovine skin – 225 g Bloom, Type B, Sigma-Aldrich*) was mixed in water at a temperature of 65 °C while stirring, to prepare 5%_w_/_w_ gelatin gels. Different fluids were injected via tattoo machine with different reciprocating frequencies of needle array (46, 60 and 72 Hz). To estimate the volume of liquid delivered inside the gel, snapshots of fluid infusion inside the gel were taken after every complete needle retraction, and then analyzed with a Matlab script to estimate the dimensions of the pattern formed by the injection.

## RESULTS AND DISCUSSION

### TATTOO INJECTION MECHANISM

Figure 2 shows schematics, and snapshots depicting the salient features of the tattooing process on the gel substrate by a 5F needle array. The puncturing process was observed to occur in four distinct phases for the first injection and for reference, we have termed them; the (*i*) piercing, (*ii*) relaxation, (*iii*) retraction and (*iv*) deposition phases sequentially. Upon actuation, the needles move downward and as they contact the substrate, the piercing phase begins. The needle array coated with a thin film of the fluid used, punctures the gel substrate at an insertion speed of *υ_i_* ∈ [0.24 0.38] m/s. The gel surface deforms in a concave shape as the needles move downwards (*figure 2(b): t = 1.5 ms*). This continues until the gel surface reaches maximum deformation, after which it begins to relax, pulling back upwards, however needle piercing continues downwards (*figure 2(b): t = 6 ms*). The needle reaches a maximum displacement corresponding to a time *t_1_* (*figure 2(b): t = 6.9 ms*) at which motion is halted before moving in the opposite direction with a retraction speed of *υ_r_* ≈ *υ_i_*.

**Figure 2:**
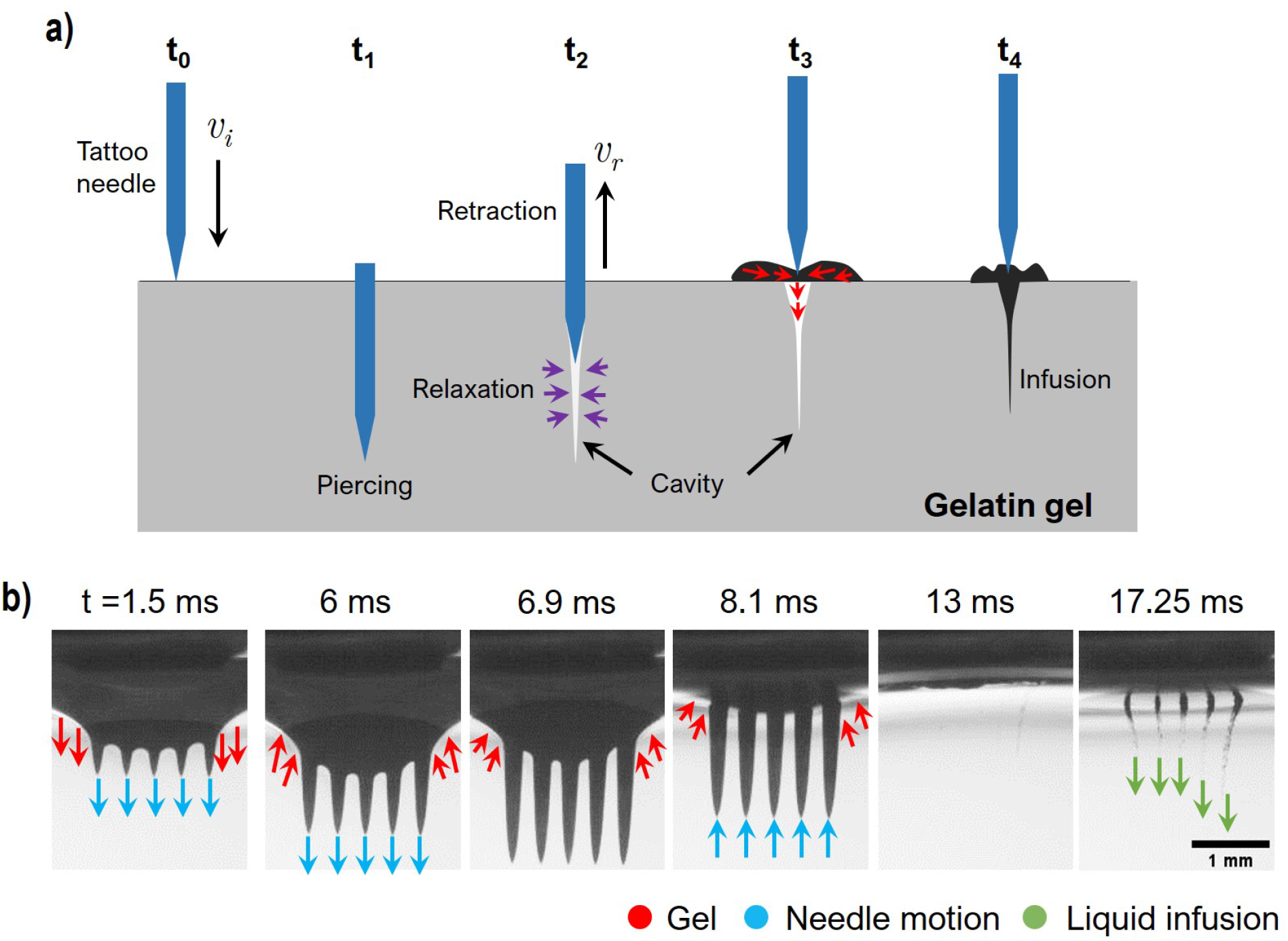
**(a)** Mechanism of fluid deposition via tattoo injection and **(b)** Snapshots showing black ink deposition for *f* = 60 Hz.

As the needles retract, cavities were formed and gel surface starts healing back to the initial shape (*figure 2(b): t = 8.1 ms*), this constitutes the retraction phase. During retraction, cavities formed decrease in size due to elastic relaxation forces in the gel matrix (*figure 2(a): t* = *t*_2_). The gel surface follows the motion of the needles as a result of adhesion forces between the gel and the needles, and was pulled upwards right before complete retraction. The surface of the gel deformed in a convex shape during the retraction of the needles from the surface (*figure 2(b): t = 13 ms*). The retraction of the needles left cavities open to the surface and to be filled by the fluid (*figure 2(a): t* = *t_3_*). At time *t_4_*, the needle has just retracted completely, leaving a cavity behind. This cavity was then filled at a rate depending on the dimensions of the cavity formed, wettability of the substrate, and the physical properties of the fluid. Finally, in the infusion phase, the needles were completely withdrawn and fluid (ink) infusion begins to occur at *t* = *t*_4_ (*figure 2(b): t = 17.25 ms*). Üpon complete needle retraction, fluid in the vicinity of the cavities left by the needles are driven in by capillary forces.

For further insight into the fluid infusion process, we performed injections with bare needles (hereafter referred to as “dry injections”) and with needles coated with the red ink. Figure 3 shows this comparison during the first injection. At the incipient stages, piercing and deforming is qualitatively similar, however distinctions become observable in the relaxation phase (*t=6.75 ms*). In case of needles coated with fluid, gel surface move upwards earlier as compared to the case of bare needles. The fluid coating minimizes the adhesion forces between the needle the gel resulting in earlier relaxation of the gel surface. It should be noted that the cavities were not visible in the case of bare needles injection and can be seen as fluid infused inside the gel in the case of needles coated with the red ink.

**Figure 3.**
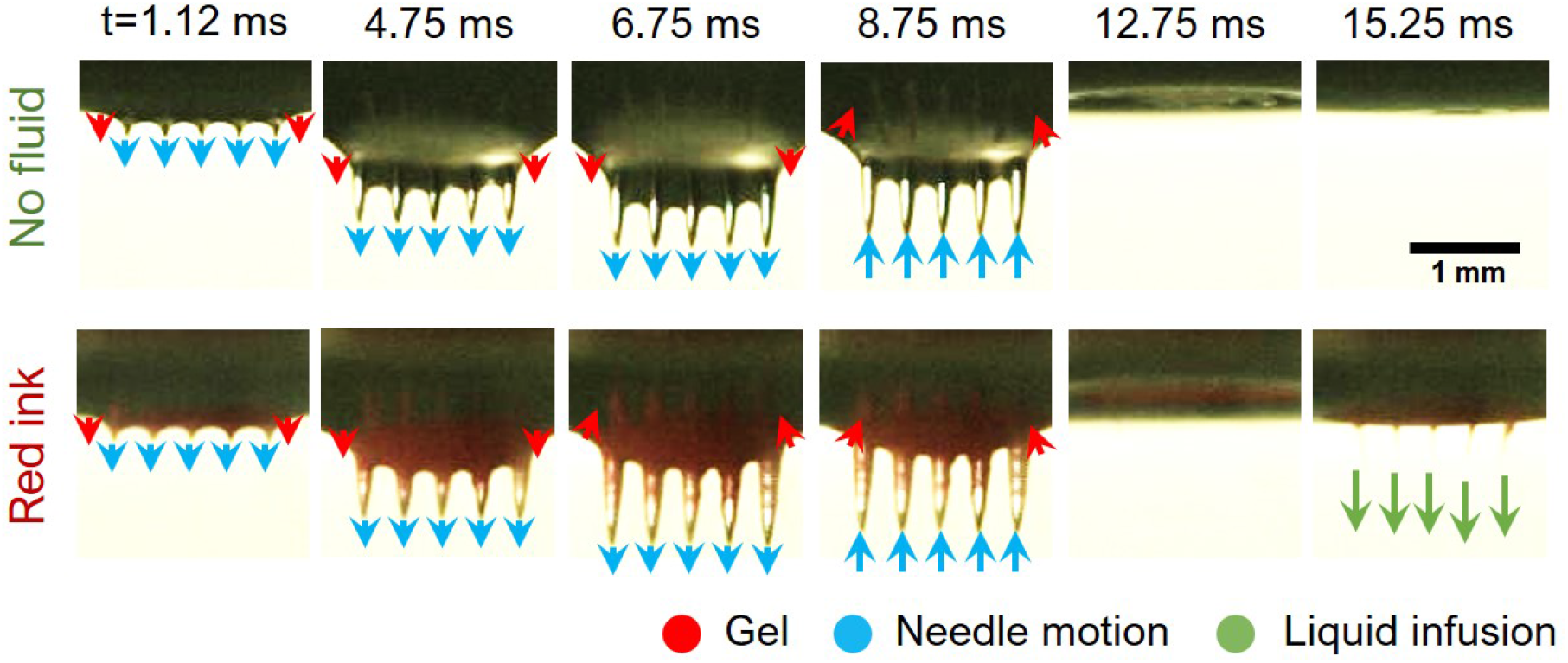
Snapshots showing the needles puncturing 5%*_w/w_* gelatin gel, with and without fluid coating.

### VISCOSITY AND RECIPROCATING FREQUENCY EFFECTS

The reciprocating frequency of the needle array was controlled by varying the applied voltage. Here we used f in a range of 46-72 Hz for infusion of two Newtonian (DI water and 80% glycerol) and two non-Newtonian fluids (red and black ink). Figure 4 shows the effect of varying f for different fluids after the first and after 50^th^ injection. The top strip of figure 4(a) shows the needles coated with the red ink at maximum penetration depth for different number of repeat injections (N). It is noteworthy that with increasing N, fluid coating visibly extends to the tip of the needles at N>5. Also, the cavities formed by the needles were observed to transit from the elastic deformation to plastic deformation. This transition can be observed with the changing shape of the cavities filled with the red ink in lower strip of figure 4(a). Thus, the amount of fluid infused (*for the red ink in this case*) increased with the increasing N (*figure 4(a)*). This phenomenon has been observed previously [20]. The projected area of the cavity filled with ink was measured from snapshots after every injection using a custom Matlab script. Assuming ink-filled cavities to have a conical shape, we estimated volume infused from the measured projected area. The volume of black ink infused in 5%*_w_/_w_* gel increased nearly linearly with increase in N for f = 60 Hz (*figure 4(b)*).

**Figure 4:**
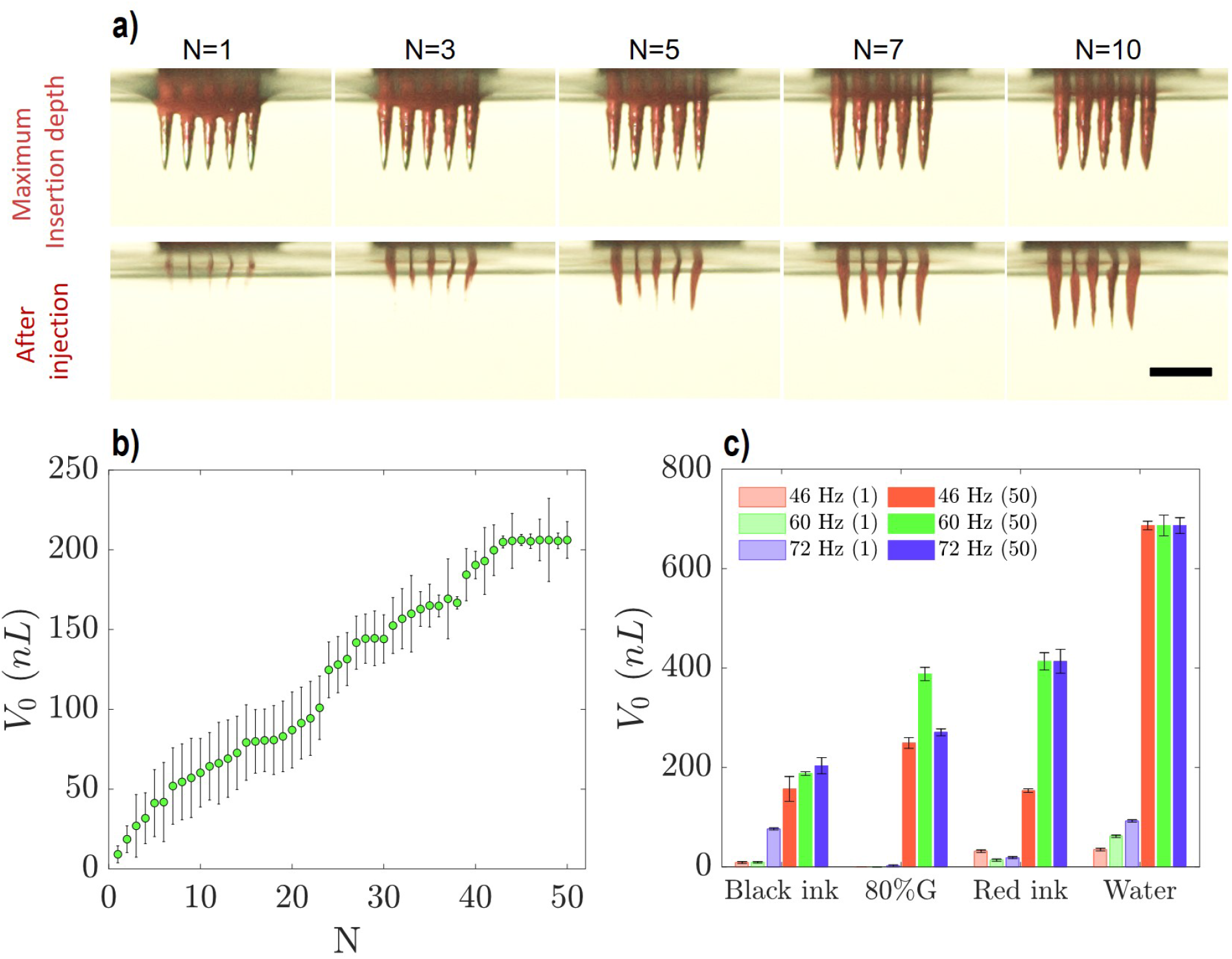
Effect of repeat injections (N) and reciprocating frequency on the infused volume and percentage delivery for 5F needle array in 5%*_w/w_* gelatin gel. **(a)** Snapshots showing needle array coated with red ink at the maximum insertion depth and after the retraction for different repeats of injections at *f*=60 *H_z_* (*Scale bar represents 1 mm*), **(b)** Volume of black ink infused with repeat injections at *f*=60 *H_z_*, **(b)** Effect of varying *f* on *V_o_* for different fluids after first (N=1) and 50^th^ injection (N=50), and (**c)** Percentage delivery of different fluids at different frequencies after 50^th^ injection.

To screen out viscosity effects, we contrast fluid deposition across all fluids for the first injection, (*N* = 1, lighter shades) and the 50^th^ injection, (*N* = 50, darker shades) as seen in figure 4(c). At *N* = 1, volume infused increased with decreasing viscosity as is evident from the higher values for black ink and water as against red ink and 80%G, a behavior that is less obvious at the lower frequencies. By the 50^th^ injection, any subtle difference between the volume infused is magnified. The Newtonian fluids still maintain a qualitatively similar behavior in volume infused, with water having a volume delivered at least 1.6 times any other fluid used (likely because of the likeness of water to the gel surface). However, the tattoo inks behave differently, more red ink is deposited into the substrate than black ink at higher *f* (an opposite of the case in the first injection). We infer that this is as a consequence of the complex properties of these fluids coming into play. Shear thinning effects may be at play seeing as an increase in *f* should correspond to an increase in shear rates. At the estimated shear rates (see *Supplementary information*) for the parameter space used in this study, both non-Newtonian fluids have been sheared to nearly the same apparent viscosity and should in theory behave the same way. It should be noted that the shearing occurs only to the fluid in contact with the moving needles. However, liquid filling the cavities after needle retraction could not have been sheared and thus, shows lower infusion for higher viscosity corresponding to low shear rates. Nonetheless, it is clear from figure 4(c) that a lower viscosity is preferable in regards to increasing delivery volume. The caveat to this premature conclusion is that our study is purely in vitro. It thus remains to be seen if this holds ex vivo.

## CONCLUSIONS AND OUTLOOK

A preliminary study of the efficacy of the tattooing process for ID drug delivery is undertaken. The major effects outlined are from the physical properties of the fluid. Shear thinning effect was limited to the fluid coated on the needle and has a minimal effect on fluid infusion due to capillary action after the needle retraction. Furthermore, multiple punctures are indeed needed to deliver more fluid into the substrate but limited by the saturation of the capacity of the cavity created by puncturing. Ultimately, our study, albeit preliminary in nature, presents an interesting baseline for any future research that may be concerned with the physics of tattooing. A deeper understanding of this process could in theory provide the basis for optimization of the tattoo procedure in any aspect that it may be used. We recommend that future mechanistic studies focus on whether our findings translate to real tissue (ex vivo, in vivo) but also to incorporate a more realistic application of tattooing whereby the needle head is translated across the surface. These aspects are the subject of our ongoing/future studies.

## AUTHOR CONTRIBUTIONS

**Conceptualization**, J.M.; **Methodology**, J.M. and I.L.; **Experiments**, I.L.; **Data Analysis**, I.L and P.R.; **Writing – Original Draft**, I.L. and P.R.; **Writing – Review and Editing**, J.M., I. L. and P.R.; **Funding Acquisition**, J.M.

## ACKNOWLEDGMENTS

J. M. acknowledges funding support through NSF (CAREER award no. 1749382).

