## Supplemental information document for "Drug delivery via tattooing: Effect of needle and fluid properties"

### Physical characterization of fluids and needle velocity measurements

Rheological profiles for black and red ink are presented in figure S1(a). A cross-model was fitted to the rheology profiles to get the apparent viscosity as: $\mu_a = \mu_\infty + (\mu_0 - \mu_\infty)/1 + (\lambda\dot{\gamma})^n$, where $\mu_\infty$ and $\mu_0$ are first and second Newtonian plateau, $\lambda$ and $n$ are constants which are obtained by the model fitting. Insertion and retraction velocities with variation in reciprocating frequency of 5F needle array is presented in figure S1(b). A linear fit yields a relation between the needle array velocity and the reciprocating frequency ($f \in [20\ 200]$) given by, $v = 0.005442f - 0.01247$. Velocity of needle array was estimated to be [0.237, 0.314, 0.379] m/s from this linear fit corresponding to frequency used in the study $f \in$ [46, 60, 72] Hz.

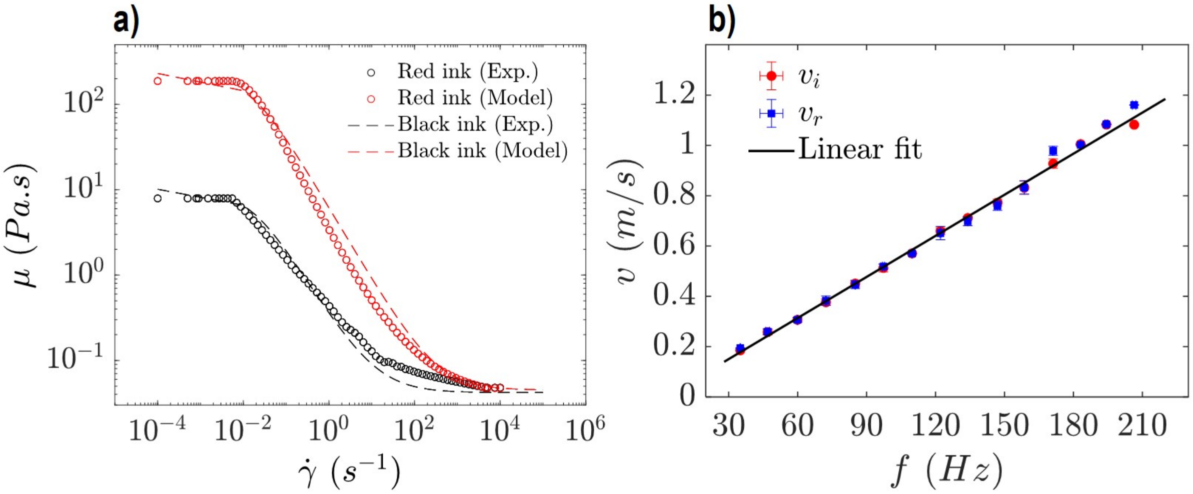


Figure S1: **(a)** Rheological measurements for different inks used and **(b)** insertion and retraction velocities of 5F needle array for different $f$ with a linear fit.

Applied shear rates ($\dot{\gamma} = 8v_i/D_{needle}$) were in a range of 5,417.1.3-8,662.9 $s^{-1}$ corresponding to $v_i \in [0.24\ 0.38]$ $m/s$ and $D_{needle}$ of 0.35 mm. Estimated apparent viscosity ($\mu_a$) corresponding to these shear rates for black and red inks were ∼0.042 $Pa.s$ and ∼0.049 $Pa.s$ respectively. Viscosity of DI water and 80% glycerol were used as $8.9 \times 10^{-4}$ $Pa.s.$ and 0.084 $Pa.s.$ respectively.
